# One experiment to rule them all? Testing multiple drivers of the temperature-size rule with nonlinear temperature increase

**DOI:** 10.1101/2020.03.16.993212

**Authors:** Andrea Tabi, Aurelie Garnier, Frank Pennekamp

## Abstract

The temperature-size rule (TSR) describes the inverse relationship between organism size and environmental temperature in uni- and multicellular species. Despite the TSR being widespread, the mechanisms for shrinking body size with warming remain elusive. Here, we experimentally test three hypotheses (differential development and growth [DDG], maintain aerobic scope and regulate oxygen supply [MASROS] and the supply-demand hypothesis [SD]) potentially explaining the TSR using the aquatic protist *Colpidium striatum* in three gradually changing and one constant temperature environment crossed with three different nutrient levels. We find that the constant and slowly warming environments show similar responses in terms of population dynamics, whereas populations with linear and fast warming quickly decline and show a stronger temperature-size response. Our analyses suggest that acclimation may have played a role in observing these differences among treatments. The SD hypothesis is most parsimonious with the data, however, neither the DDG nor the MASROS hypothesis can be firmly dismissed. We conclude that the TSR is driven by multiple ecological and acclimatory responses and hence multicausal.

**Author statement:** A.T. designed and led the experiment, and A.T. and A.G. performed the sampling. A.T. and F.P. analyzed and interpreted the data. All authors contributed to the writing of the manuscript.

## 1 Introduction

Temperature and body size jointly influence biological rates from the individual to the ecosystem level (Brown et al. 2004). But temperature and size are not independent. About 80% of ectotherms are smaller when grown in warmer environments (Atkinson 1994). This pattern is so widespread that is has been named the temperature-size rule (TSR), describing the negative correlation between body size and environmental temperature observed both in multi- and unicellular organisms (Atkinson 1994, Atkinson et al. 2003, Forster and Hirst 2012, Forster et al. 2013). Despite the TSR being widespread and having important consequences for the structure and functioning of communities, the mechanistic drivers of the TSR remain elusive.

Multiple hypotheses have been proposed and tested to explain the TSR (Figure 1A). The differential development and growth (DDG) hypothesis states that differences in thermal sensitivities of growth and development rates of organismal ontogeny explain the TSR, i.e. temperature changes the growth/reproduction trade-off (Forster et al. 2011). The DDG hypothesis assumes that the rates of cell division and differentiation are more sensitive to temperature than cell growth and hence increased temperature would result in smaller organisms (van der Have and de Jong 1996). If, however, cell growth would be more strongly affected than development, larger sized individuals would result. Such a reversed temperature-size rule, where the cell sizes are larger at higher temperatures, has been observed as well (Zuo et al. 2012). For unicellular organisms, Forster and colleagues (2011) suggested that the ratio between development and growth rates should be similar between temperatures due to their binary mode of division. Variation in this ratio would hence only occur during acclimation to novel temperature regimes and indicated by a temporary change in the ratio of mother and daughter cell sizes (Forster et al. 2013).

**Figure 1:**
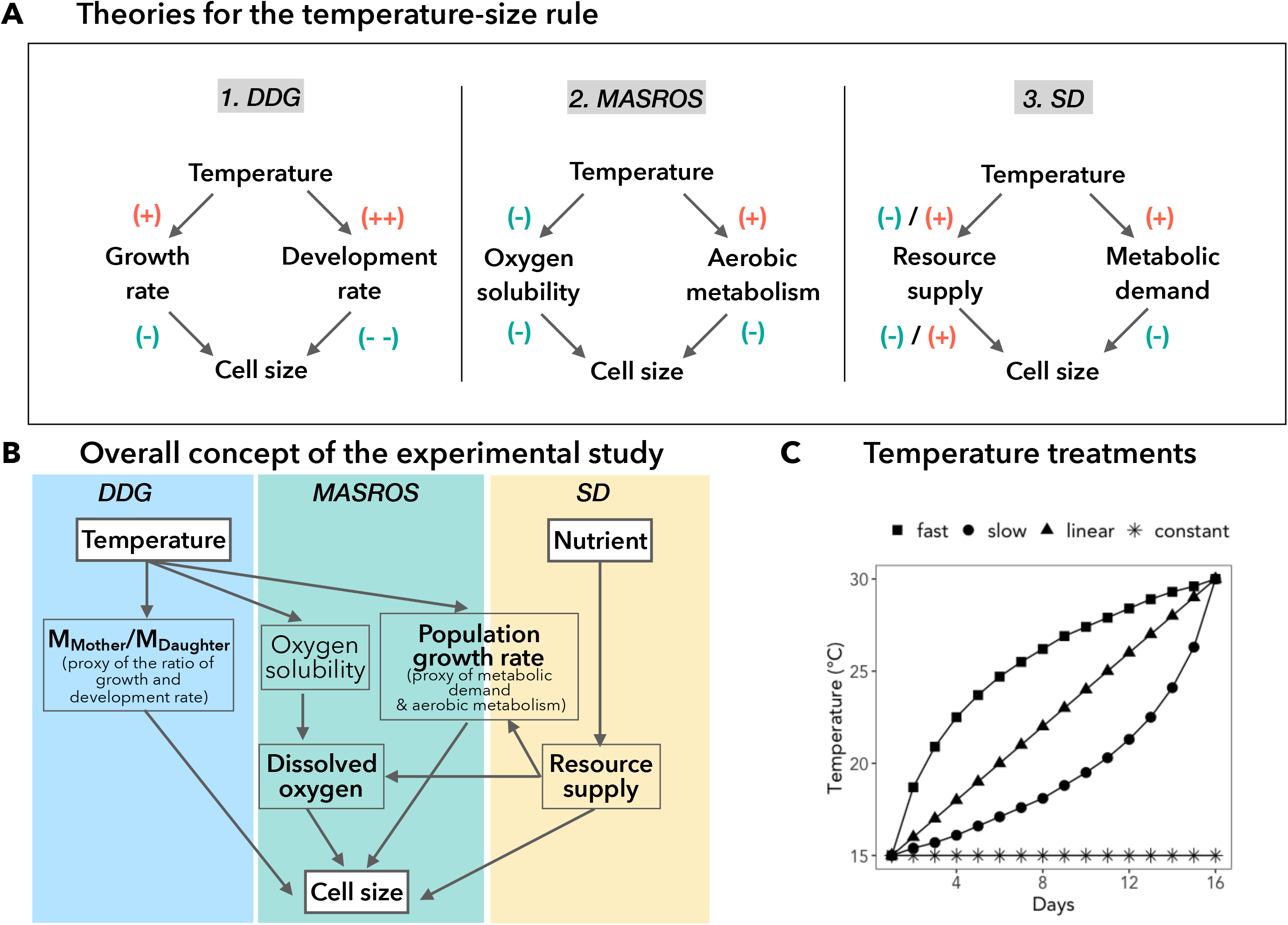
Theoretical and conceptual background. **(A)** Three hypotheses have been proposed to explain the mechanistic drivers of the TSR. (1) The differential development and growth (DDG) hypothesis states that in warm environment an organism has less time to grow, because warming accelerates the rate of development (++) faster than that of growth (+), which results in a smaller cell size. However, a reversed TSR has been observed in some cases, which might indicate that growth has higher sensitivity relative to development. (2) Elevated temperature affects oxygen availability through decreasing oxygen solubility (-) in the water and also through increasing aerobic metabolism (+), which both bring about smaller cell size. (3) Increased temperature also has a positive effect on the metabolic rate (+), which increases resource demand of the organism relative to the available supply. Resource supply can either be positively or negatively (+/-) affected by temperature. This mismatch of demand and supply then leads to a smaller cell size (supply-demand model). **(B)** In this study we jointly test empirically these three hypotheses, in order to identify the major drivers of TSR and their interconnections. Furthermore, we also account for the effect of nutrient enrichment on cell size. **(C)** The four different temperature treatments applied: constant 15°C, linear, slow and fast rising temperature treatment between 15-30°C. The fast and slow rising temperature treatments have opposite shapes. Not shown is the nutrient treatment (three levels) fully crossed with the temperature treatment.

Despite direct temperature effects on the physiology of individuals, it is also possible that temperature affects size indirectly, that is, mediated by another environmental factor that is influenced by temperature. The MASROS (i.e. maintain aerobic scope and regulate oxygen supply) hypothesis proposes that organisms change their body size to regulate the oxygen demand of their tissues (Atkinson et al. 2006) (Figure 1A). At higher temperatures organisms have increased metabolic oxygen demand. At the same time, the oxygen solubility decreases in warmer waters, hence limiting the oxygen diffusion to the cell surface. Being smaller becomes more beneficial due to a higher surface to volume ratio (Atkinson et al. 2006). The MASROS hypothesis is expected to be of great importance in aquatic environments, where the oxygen solubility of water is strongly influenced by temperature (Forster et al. 2012).

Finally, size change with temperature could be driven by the need of organisms to balance their bodily demands for growth with the supply of resources from the environment (the supply–demand model, SD) (DeLong 2012). The supply-demand hypothesis posits that organisms balance their demand and the supply at an optimal body size for a given environment. According to this hypothesis, the TSR is consistent with the idea of balancing increasing metabolic demand due to temperature with a constant or decreasing environmental supply. The SD hypothesis provides an explanation for possible reversals of the TSR, as well as oscillations in cell size (DeLong et al. 2017).

While the TSR hypotheses only refer to changes in cell size, high temperatures also affect other morphogenetic processes such as cell shape. For example, at elevated temperature *Tetrahymena thermophila* showed oscillations between more oblong and rounder shaped cells which negatively correlated with cell size (DeLong et al. 2017). Another study found that cell shape differed significantly with increasing temperature, linking changes in shape and size together, and cultures at highest temperatures contained teratogenic shapes (Neustupa et al. 2008).

Although all TSR hypotheses have been empirically tested, there is no consensus about the most likely mechanistic drivers. This is because previous experiments mostly tested a single hypothesis at a time (see Table S1 in Supplementary Information). However, these hypotheses are not all mutually exclusive, but instead might complement each other, i.e. the drivers behind the TSR could be a combination of multiple effects, that is, they are multicausal. To our knowledge, only DeLong et al. (2017) discussed the three hypotheses, however, relying only on indirect evidence for the MASROS and DDG hypotheses and hence not providing a conclusive answer. Therefore, simultaneously evaluating all three hypotheses experimentally on a single dataset is needed.

Previous experiments have studied the TSR without allowing for acclimation. The majority of experiments manipulated the temperature with an abrupt change compared to the control (see Table S1). This is in contrast to the continuous and gradual temperature change that organisms experience in natural conditions. It is therefore possible that the temperature-size response is exaggerated due to the abrupt exposure of organisms to large temperature changes, whereas the response may be ameliorated by acclimation of individuals to more gradual temperature change. To understand the role of acclimation, we need studies that evaluate the effect of gradually changing temperatures on the TSR. An experimental design that can inform about the acclimation process is the comparison between constant and a rising temperature (e.g., Mitz et al. 2019, Peck et al. 2013).

Here, we aim to understand the relationship between cell size and gradually increasing temperature using the ciliate *Colpidium striatum* as our study organism. We experimentally test all three hypotheses regarding the mechanisms of TSR in aquatic microorganisms (Figure 1B). To understand the role of acclimation, we used one constant and three gradually increasing temperature treatments, where the rate of temperature change is either constant through the experiment (i.e., *linear*), a fast rise during the growth phase (i.e., *fast*) or a fast rise during the stationary phase (i.e., *slow*, see figure 1C for an illustration). Responses to the different temperature can hence inform about the organism’s ability to acclimate to warming on different time scales (i.e. intra- vs intergenerational).

We hypothesize that if temperature changes faster than the organism can acclimate, decreasing abundance and/or extinctions will be observed. On the contrary, if organism can keep up with the temperature change, we expect to see TSR, i.e. shrinking cell size and dynamics similar to the constant temperature treatment. At high speed of increase into sublethal temperatures, we expect to see various responses such as vegetative enlargement to increase viability without inducing sexual reproduction (Atkinson et al. 2003), whereas in the optimal thermal range an inverse relationship between size and temperature is expected. We predict a decoupling of cell mother and daughter cell size ratios in all but the constant temperature treatment. We also predict an increased growth rate with temperature increase, whereas resource supply is expected to remain constant.

## 2 Materials and methods

### 2.1 Experimental design

Microcosms were loosely capped 250 mL glass jars containing 100 mL of medium (0.28, 0.56 or 1.12 g/L of Carolina Biological Supplies Protist Pellet in Chalkley’s medium) (Altermatt et al. 2015) with two autoclaved wheat seeds added for the slow release of carbon and nutrients.

A mix of bacteria (*Bacillus subtilis, Serratia fonticola* and *Brevibacillus brevis*) forms the basal food supply of the bacterivorous protist *Colpidium striatum*. We inoculated bacteria in the medium 2 days prior to the experiment. We then added 1 mL *Colpidium* drawn from stock cultures kept at 15 °C, to obtain an initial concentration of 3 individuals per mL and randomly assigned microcosms to temperature treatments.

The experiment has a full-factorial design of four levels of temperature treatment (constant 15°C, three rising temperature between 15-30°C, see Figure 1C) and three levels of nutrient concentration (low = 0.28, medium = 0.56, and high = 1.12 g Protist Pellet medium per liter). We choose constant 15°C as the reference temperature as this is representative for the temperature *Colpidium striatum* experiences in natural freshwater environments. All three rising temperature treatments start at 15°C and end at 30°C. The linear speed is +1°C per day, while the fast rising temperature increases at the beginning of the experiment faster then it slows down and the slow rising temperature treatment goes in reversed fashion. The selected temperature treatments reflect the predicted per generation rates of increase over the next decades (IPCC 2007). Furthermore, this temperature range is suitable for studying dynamical changes based on the thermal response of the selected species. Based on previous experiments, the abundance and growth rate of *Colpidium striatum* increases up to 17°C and then decreases in the 17-30°C temperature range (Jiang and Morin 2004, Plebani 2015). We expect that the bacteria can deal with environmental change better than protists due to faster reproduction and a wide thermal range (from 10 to 40°C), hence maximum density should be comparable across temperature treatments but would differ across nutrient availability. Temperature was manipulated by use of four incubators. This design results in 12 treatment combinations, with five replicates each, and 60 experimental units in total.

During the experiment, 5 mL of medium (of a total of 100 mL medium) was taken every day and replaced with 5 mL of sterile media. 1 mL of the sample was transferred to a counting chamber and filmed under a stereo microscope with a 25× magnification mounted digital CMOS camera (Hamamatsu Orca C11440; Hamamatsu Photonics, Japan) to take a 5s video (25 frames per second). The following properties were measured with the R package BEMOVI (Pennekamp et al. 2015): individual morphology (cell size and shape), and population size (individuals per mL). Furthermore, samples were processed to measure bacterial abundance using flow cytometry with an Accuri C6 flow cytometer and dissolved oxygen (DO) concentration with oxygen sensors (from PreSens - Precision Sensing GmbH); for details refer to Tabi et al. (2019).

### 2.2 Response and predictor variables to understand the TSR

Cell size was estimated for each individual using length and width information assuming an ellipsoid form (Laakso et al. 2003): 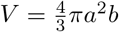, where *a* is the radius of width and *b* is the radius of length. Then, we calculated the cell size as the median cell volume (MCV) for each day in each population. The mother and daughter cells were estimated as the upper and lower 20% of cell size distribution for each day. Cell shape was measured by the aspect ratio (cell length / cell width) of each cell. The aspect ratio closer to 1 represents a rounder form, whereas >1 means a more oblong cell shape.

The growth rates for each population were estimated by fitting logistic growth model with changing intrinsic growth rates to the time series of *Colpidium*’s density (Werker and Jaggard 1997):

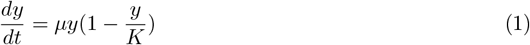

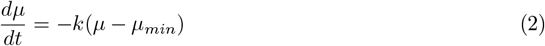

where *μ* is the growth rate, *K* is the carrying capacity and *k* is a constant which determines the speed with which the initial intrinsic growth rate approaches the final minimum intrinsic growth rate, *μ_min_*. The carrying capacities were calculated as the maximum density of the time series of each population. The growth rate was used as the demand proxy to test the SD hypothesis. Bacterial abundance represented the supply side in the SD model. The dissolved oxygen concentration of the medium was used for testing the MASROS hypothesis.

### 2.3 Data analyses

#### 2.3.1 Mother to daughter cell size ratio

To test the DDG, we relied on the equation by van der Have and de Jong (1996), i.e. 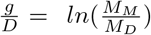, where *g* is the mass-specific growth rate of the individual (day^−1^), *D* is the development rate (day^−1^, that is, 1/doubling time), *M_M_* is the mass of the mother cells and *M_D_* is the mass of the daughter cell. Deviations from the expected ratio of ln(2) indicate acclimation (Forster et al. 2013).

#### 2.3.2 Linear model analysis

In order to understand the relationship between cell size and population abundance, temperature, time and nutrients, we used a general linear model:

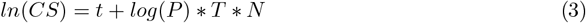

where *CS* is the cell size measured as the median cell volume (MCV), *t* is time (days), *P* is the cell density (cells mL^−1^), *T* is temperature (°C) and *N* is the nutrient concentration (three levels: low, medium, high). Cell size and density were log-transformed to meet assumptions of homoscedastic residuals. For the constant temperature treatment, we added a small amount of noise to the constant 15°C time series to be able to estimate the temperature term. Noise was modelled as a Gaussian distribution (mean = 0, SD = 0.5), representative for the small temperature fluctuations which naturally occur in the incubators. For the linear treatment, we dropped the day term because it is perfectly correlated with the temperature term, and hence it is not possible to estimate both.

#### 2.3.3 Structural equation modeling

We fitted structural equation models (SEM) to the data to understand the drivers of changes in median cell volume (Grace et al. 2010) according to the SD and MASROS hypothesis. The DDG hypothesis was tested in a separate analysis as the ratio does not directly predict the change in cell size. Relevant predictor were the time series of temperature, the nutrient availability (three levels: low, medium, high), the mean cell density (individuals per mL), the bacterial abundance (cells per mL as a supply proxy), the growth rates (as a demand proxy), as well as the dissolved oxygen content measured in the medium. We started with an initial SEM including the causal drivers specified in Figure 1B and fitted the model simultaneously to the four temperature treatments. We then evaluated model fit with the Chi square statistic and the comparative fit index (CFI) to find a model that captures the covariance structure of the data. We updated the model until all relevant links were included. We then simplified the model by removing links which were included in the initial model but had relatively low explanatory power with AIC-based model comparisons. Finally, we tested whether coefficients were different across groups, by comparing models where certain path coefficients were forced to be the same, against freely estimated coefficients. Model comparison was based on the AIC. All structural equation models were fitted with the lavaan package (Rosseel 2012) in R.

## 3 Results

### 3.1 Cell size, shape, and population abundance through time

In the constant treatment population dynamics closely followed logistic growth, i.e. after an initial growth phase, carrying capacity was reached and maintained till the end of the experiment. The slow warming treatment showed remarkably similar dynamics, despite the fast increase in temperature at the end of the experiment. The fast warming treatment showed initial growth and a short stable phase, whose length depended on the nutrient treatment. The linear treatment showed dynamics intermediate to the slow and the fast increase (Figure 2).

**Figure 2:**
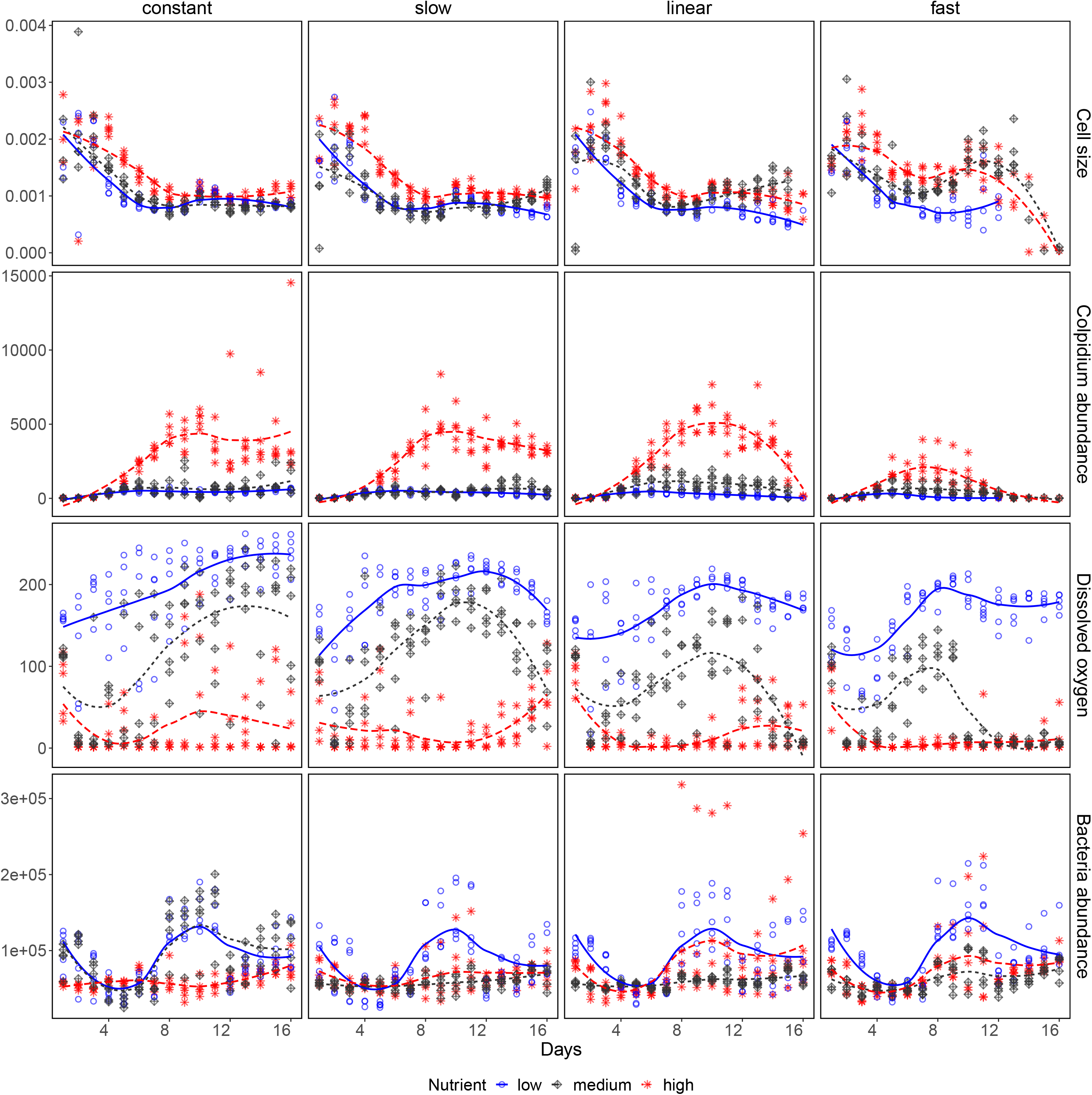
Time series of *Colpidium striatum* cell size and abundance, dissolved oxygen content and bacteria abundance by temperature treatment. The population abundance of *Colpidium striatum* and bacteria is measured as individuals mL^−1^. Different nutrient treatments are colour coded. Lines (spline fits) are used to highlight the trends in the data.

Cell size decreased with time in the constant, slow and fast warming temperature treatments (Figure 2, Figure 3, Tables S2-S3 in Supplementary Information). The temperature effect was not significant for all but the fast treatment, where it was positive, but note that temperature interacted with other factors rendering the interpretation of the main effect rather difficult. The effect of enrichment was only visible in the linear and fast temperature treatments, where it negatively affected cell size in the medium enrichment case, and positively in the high enrichment case, respectively (Figure 3). Density and temperature interacted in the fast change treatment only, whereas both nutrient enrichment interacted with temperature in a positive way, hence alleviating the negative effect of temperature, for all but the fast treatment. Density interacted with enrichment in a negative fashion in the fast treatment only, while the three-way interaction was positive only for the fast treatment (Table S3).

**Figure 3:**
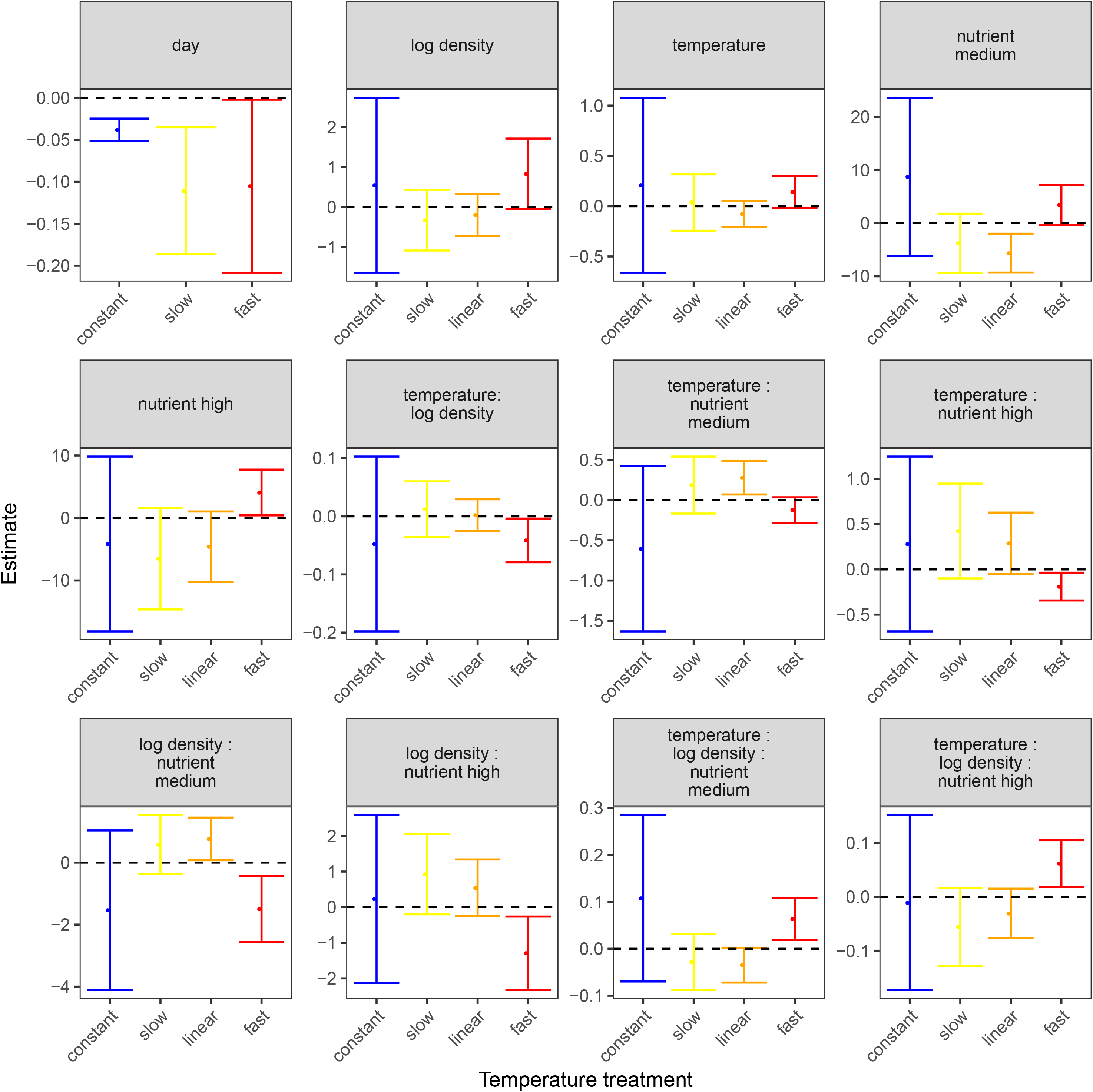
Linear model parameters estimating effects of time (in days), density, temperature and nutrient enrichment on cell size. The dashed lines indicate zero. Error bars show 95% confidence intervals for the estimates.

To better understand the consequences of interactions among factors on cell size, we show plots of the estimated effects for the four treatments in the Supplement (Figure S3-S6). The interaction plots suggest that slowly increasing temperature had a positive effect on cell size in the medium and high nutrient enrichment, but the temperature effect reverses to be negative in the low enrichment treatments, when warming is linear or fast. Increasing density reinforced the temperature effect across all increasing temperature treatments.

Cell shape also varied with time (Figure S2). The changes in cell shape were generally negatively correlated with the cell size over time (see Table S5 in Supplementary Information). In constant 15°C, both the cell size and shape stabilized showing a strong negative correlation across all nutrient enrichment. In fast and linear warming combined with low nutrient enrichment, no strong pattern was observed. In both cases, the aspect ratio changed from rounder to more oblong and then back to rounder, while cell size decreased. Variation in aspect ratio was highest in the fast treatment with medium and high nutrient enrichment, which also mirrored the changes in cell sizes. In linear temperature treatment, after reaching a more oblong shape, cell shapes returned to rounder form at the end of the experiment across all nutrient enrichment.

### 3.2 Understanding the drivers of the temperature-size rule

The log ratio of mother and daughter cells showed no decoupling of mother and daughter sizes (Figure S1), even after removing the effect of population abundance on cell sizes. Instead, the log ratio of mother and daughter cells did not change over time but fluctuated around the expected value of ln(2) in constant and all rising temperatures, regardless of nutrient treatment.

The final structural equation model was a good fit to the data (comparative fit index: 0.999, test statistic 15.947, df = 14, p = 0.317) and explained between 30% and 45% of the variation observed in cell size (see Table 1). Variation in cell size was best explained in the constant and slow warming treatments. Across treatments, variation in oxygen content of the medium was best explained (*R*^2^: 0.56 - 0.63), whereas abundance of bacteria had the lowest explanatory power (*R*^2^: 0.08 - 0.11).

**Table 1:** Coefficients of determination (R2) for the five endogenous variables in the SEM.

| variable | fast | linear | slow | constant |
| --- | --- | --- | --- | --- |
| Cell size | 0.3 | 0.31 | 0.43 | 0.45 |
| Dissolved Oxygen content of medium | 0.63 | 0.61 | 0.6 | 0.56 |
| Consumer abundance | 0.22 | 0.44 | 0.5 | 0.43 |
| Bacteria abundance | 0.11 | 0.11 | 0.08 | 0.08 |
| Growth rate | 0.77 | 0.66 | 0.37 | 0.11 |

The SEM indicated that cell size was impacted directly and indirectly by temperature (see Figure 4 and Table S4). The direct effect of temperature on cell size (MCV) was negative in the fast and linear warming treatments. However, temperature also affected cell size indirectly, mediated by the dissolved oxygen content of the medium. Temperature increased the dissolved oxygen content in the linear and slow treatments, whereas it had no effect in the other treatments. Dissolved oxygen content itself affected cell size in a consistently negative fashion across treatments, meaning higher oxygen content led to lower cell size. Temperature had a negative effect on growth rate for all but the constant treatment, resulting in more strongly decreased growth rates with increasing speed of warming. All increasing temperature treatments had a positive effect on bacteria abundance, whereas the consumer abundance remained relatively constant across warming treatments. Higher bacteria abundance did not directly translate into larger cell size. Surprisingly, the direct effect of growth rate on cell size was positive in the constant and slow treatment, opposite to the expectation for a demand proxy. Consumer abundance had the expected negative effect on cell size, for all but the fast treatment where it was positive.

**Figure 4:**
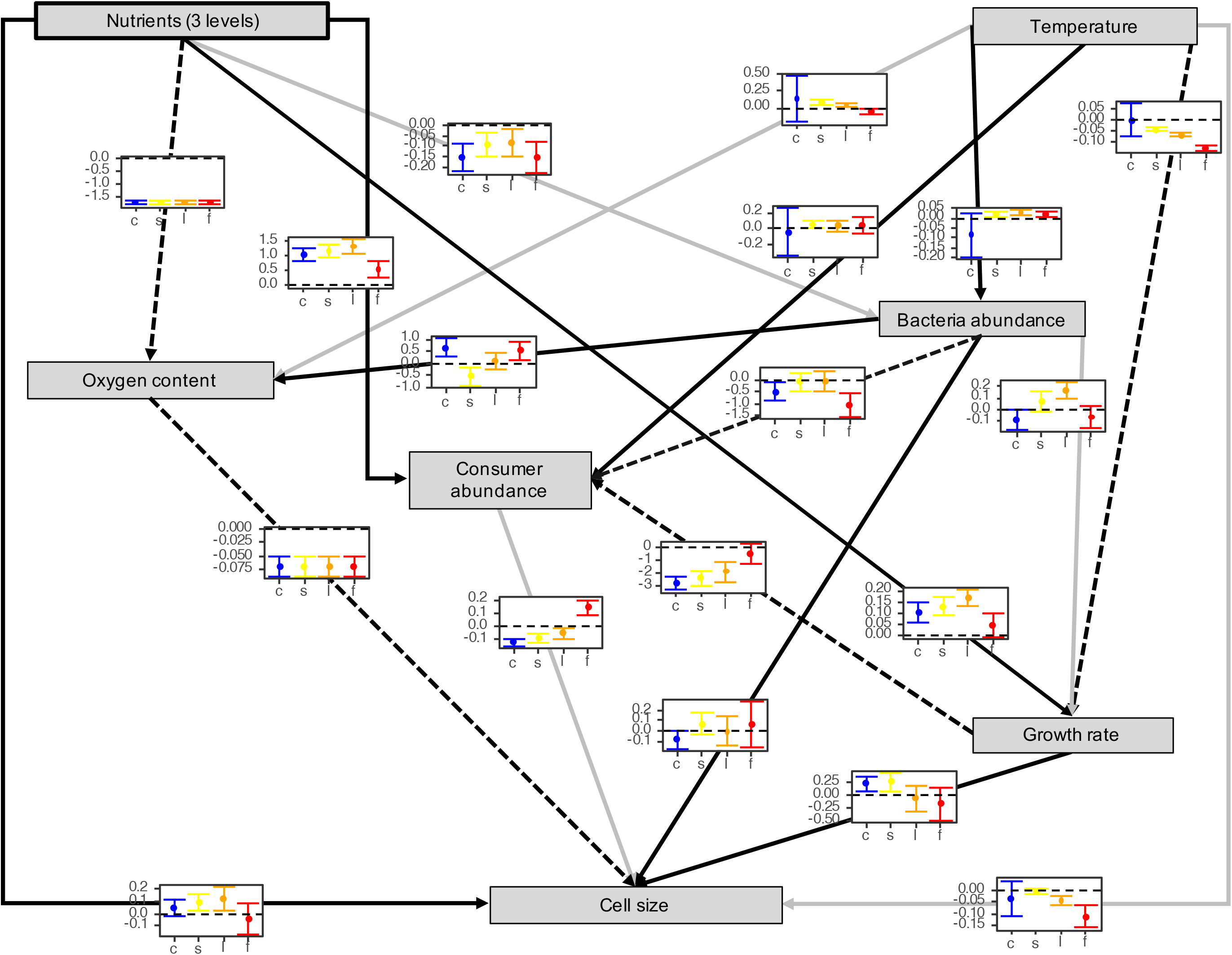
The causal network based on structural equations. The path coefficients are shown in the small inset plots, with the different temperature regimes (groups) shown with different colours (constant = blue, slow = yellow, linear = orange, fast = red). Dashed arrows indicate a predominantly negative effects, while solid arrows indicate a positive effects across temperature change. Grey arrows indicate a switch of sign in the coefficients across treatments.

Nutrients had a positive effect on consumer abundance, but a negative effect on bacteria abundance, potentially indicating a trophic cascade. This was also reflected in higher growth rates in all but the fast warming treatment. Higher nutrient concentrations nevertheless only translated into larger cells in the slow and linear treatments.

## 4 Discussion

The temperature-size rule is a widely observed phenomenon, however, the mechanisms behind the TSR remain elusive. Here, we provided the first comprehensive analysis integrating components of different theoretical frameworks on the same experimental dataset to pinpoint the mechanisms of size changes in an unicellular model organism.

We observed substantially different dynamics among the rising temperature treatments, giving an indication of the time lag needed for acclimation. As for our hypothesis regarding the temperature effects in intra- vs intergenerational time scale, we showed that if the warming occurred at higher speed affecting the generation time during the growth phase, it led to extinctions. Moreover, cell sizes and shapes showed the strongest response in the fast change treatment, supporting our hypothesis. Interestingly, the slow temperature increase and the constant 15^°^C temperature affected *Colpidium* in a very similar fashion, which suggests that cells were able to acclimate to the slowly warming environment.

### 4.1 Evidence for different hypotheses

Our analysis of the log ratios of mother and daughter cells did not show any sign of decoupling across the four treatments. Rather mother-to-daughter cell size ratios fluctuated around the expected ratio of *ln*(2), regardless of the nutrient enrichment. In a previous study with a unicellular protist, support was found for the DDG hypothesis the during acclimation phase, which took 70h and 120h for mother and daughter cells respectively (Forster et al. 2013). However, organisms were exposed to different constant temperatures, unlike the gradually changing environmental temperatures as in our study. It is possible that acclimation to gradually changing temperature masked the pronounced decoupling visible when cells were exposed immediately to higher temperatures. Nevertheless, we would have expected to see a decoupling at least in the linear and fast changing temperature treatments. Therefore, we cannot fully exclude or support the DDG hypothesis.

While the pathway between temperature and dissolved oxygen content in the medium was supported by the SEM, the dissolved oxygen negatively affected cell sizes, i.e. higher dissolved oxygen content led to smaller cell size. This response was consistent across the four temperature treatments. *Colpidium* has tricarboxylic acid cycle allowing the synthesis of proteins in anaerobic conditions (Seaman 1949), however, in the presence of bacteria its oxygen uptake can be limited. Therefore the MASROS hypothesis might have a low importance in explaining shrinking cell sizes, in line with findings by Forster and Hirst (2012), where an effect was only observed for multicellular organisms. The negative effect of dissolved oxygen on cell sizes can be an indirect effect of increasing metabolism of the protists which resulted in higher consumption of bacteria and therefore lower level of dissolved oxygen in the medium. Furthermore, the dissolved oxygen level showed no consistent pattern with either the bacteria or the the protist abundance and also cannot explain the decreasing cell sizes with temperature in our study. A potential explanation for not seeing any effect may be that *Colpidium* is only limited at very low dissolved oxygen concentrations, which we did not reach in this experiment. Therefore, we cannot fully exclude or support the MASROS hypothesis.

The supply-demand model predicts changes in cell size as a function of temperature-dependent demand and supply (DeLong 2012). Higher temperature will generally lead to higher metabolic rates and hence demand. Temperature may also affect the supply of resources. We found that temperature had a negative effect on growth rate, i.e. quickly increasing temperatures decreased growth rates more than linearly or slowly increasing temperatures. Regarding supply, temperature had a positive effect on bacteria abundance, with increasing abundance in the slow compared with the fast treatments. As expected, higher bacteria abundance (supply) had a positive effect on cell size, however, the effect was only significant in the fast treatment. The effect of temperature on the balance of supply and demand determines whether cell size should increase or decrease. Given that demand was higher in slow and linear treatments but bacteria abundance was also higher in these treatments one could expect these drivers to cancel out. Hence, although our results are consistent with the mechanisms of the SD model, we would expect to see little effect in our study. Previous experiments using the SD model found evidence for the balance between supply and demand explaining size variation in protists (DeLong 2012), including possible reversals of the temperature-size rule (DeLong et al. 2017)

It is also possible that multiple mechanisms are relevant at different time scales. To estimate the TSR in unicellulars, cells are usually measured during the stationary phase: once the cell size was acclimated and after few generations (Montagnes and Franklin 2001, Forster et al. 2013). However, within the growth phase of a single population, cell size also varies (e.g., Figure 3 in Forster et al. 2013). With environmental conditions similar to what a cell previously experienced, a new population showed higher cell size at low density, and then a decrease in size until reaching the carrying capacity. Therefore it is possible that the mechanisms explaining the TSR act at different temporal scales, depending on the supply constraints: during the growth phase, when resources are not limiting, the increased metabolic rate (and growth rate) may be most important. In contrast, when the population reaches carrying capacity (i.e., a density-dependent parameter which balances the density, the cell size and the resource use), the SD model seems to fit the observations best (DeLong et al. 2017).

In summary, our results are consistent with the mechanisms of the SD model, while the MASROS and DDG hypotheses are neither ruled out nor supported. Instead, other factors like the density of the consumer mediate the effect of temperature in our study. The direct effect of temperature on cell size also hints at other, still unidentified factors being important.

### 4.2 Acclimation to gradual temperature change

Previous attempts to study how cell size is affected by temperature have relied on designs where cells were immediately exposed to higher temperatures and the response compared to cell size in control temperatures to which cells were adapted (see examples in Table S1). Our design allows us to investigate the dynamic acclimation process as we exposed cells to gradually changing temperature regimes with a faster temperature increase occurring during either the population growth phase or the stationary phase (i.e. fast and slow treatment respectively). Our results indicate that linear and fast temperature increase leads to the occurrence of frequent interactions among temperature, density and nutrients potentially suggesting that cells were unable to adapt to the fast temperature increase, while simultaneous acclimation processes in the slow and linear treatment lead to additive processes unfold. Investigating the three-way interactions revealed that nutrient enrichment buffers some of the adverse temperature effects, but as soon as temperature change accelerates (linear and fast change treatment), a negative temperature-size effect was observed. We interpret this as a signal of the acclimation process unfolding too slowly in the linear and fast change treatments, resulting in the population collapse observed as well as the negative temperature-size rule. Indeed, in our study, *Colpidium* experienced daily a temperature change (Figure 1C), whereas observed an acclimation period between 70.5 and 120h for *Cyclidium glaucoma* (Forster et al. 2013).

### 4.3 Limitations

Our study is the first to take an integrative approach which compares the ability of different frameworks to understand the effect of temperature on cell size, combining distinct physiological pathways like oxygen consumption and the balance of supply and demand. The structural equation models allow us to compare the relative size of effects and a causal interpretation given the experimental nature. However, not all of the key variables could be manipulated, hence some of the estimated effects have to be evaluated with caution. Foremost, we only measured oxygen availability in the experimental cultures, which can co-vary with other variables such as bacterial density. While we controlled for some of these effects, our approach remains partly observational and hence future experiments should evaluate the oxygen effect by manipulating oxygen availability. The previous comment also applies to other variables that co-vary with the temperature change, such as time and density, which are difficult to disentangle due to the inherent covariation due to the growth process. While we integrate three main hypothesis regarding causal drivers of the temperature-size rule, our experiment cannot account for all possible hypotheses regarding temperature-size responses. For example, lacking measurements of DNA content, we were not able to test whether temperature induces changes at the the genomic level, which trigger subsequent changes in cell size to maintain the genome size to cell volume ratio (Hessen et al. 2013).

### 4.4 Conclusions

Our study revealed the multicausal nature of the temperature-size rule in gradually warming environments. The SEM showed that the SD hypothesis is most consistent with the observed changes in cell size for our experimental system, but the role of maintaining aerobic scope and regulating oxygen supply as well as differential development and growth cannot be firmly dismissed. Our analyses also highlight the role of acclimation and nutrient availability in mitigating the effects of temperature on organism size. How these factors act in concert needs to be considered to foresee the effects of warming on organism size in natural environments.

## 5 Acknowledgements

AT and FP were financially supported by Swiss National Science Foundation Grant 31003A_159498. AT was also supported by the National Research, Development and Innovation Office – NKFIH, grant GINOP-2.3.2-15-2016-00057.

