## Supplementary material for "One experiment to rule them all? Testing multiple drivers of the temperature-size rule with nonlinear temperature increase"

### 1 Supplementary tables

| **Authors** | **Year** | **Study system** | **Treatment** | **Theory tested** | | |
|---|---|---|---|---|---|---|
| | | | | DDG | MASROS | SD |
| Forster et al. | 2011 | meta-analysis of various systems | various constant temperatures used across studies | X | | |
| DeLong | 2012 | *Actinosphaerium* | three constant temperatures (20, 25, 28°C) | | | X |
| Forster et al. | 2013 | *Cyclidium glaucoma* | constant temperature | X | | |
| Huete-Stauffer et al. | 2016 | bacteria | daily fluctuations with three regimes (ambient, warming (+3°C) and cooling (-3°C)) | X | | |
| Walczyńska et al. | 2016 | *Coleps hirtus* | six constant temperatures (10, 15, 20, 25, 30, 35°C) | X | | |
| DeLong et al. | 2017 | *Tetrahymena thermophila* | three constant temperatures (20, 26, 32°C) | X | X | X |
| Skau et al. | 2017 | *Emiliania huxleyi*, *Chrysochromulina rotalis* and *Prymnesium polylepis* | two constant temperatures (13°and 19°C) | | | (X) |

Table S1: **Literature overview of empirical studies on TSR with an experimental approach on unicellular organisms.** DDG = Differential development and growth, MASROS = maintain metabolic scope and regulate oxygen, SD = supply-demand model. The study by Skau and colleagues did not explicitly test the SD model, but allows for the interpretation regarding the mechanisms of the SD model (indicated by (X)).

Table S2

| | *Dependent variable:* | |
|---|---|---|
| | log(MCV) | |
| | Constant | Slow |
| | (1) | (2) |
| temperature | 0.207 (0.441) | 0.036 (0.142) |
| log_density | 0.544 (1.108) | −0.325 (0.385) |
| nutrientmedium | 8.690 (7.549) | −3.782 (2.823) |
| nutrienthigh | −4.166 (7.088) | −6.512 (4.120) |
| days | −0.038*** (0.007) | −0.111*** (0.038) |
| temperature:log_density | −0.048 (0.076) | 0.012 (0.024) |
| temperature:nutrientmedium | −0.606 (0.521) | 0.186 (0.180) |
| temperature:nutrienthigh | 0.281 (0.490) | 0.424 (0.266) |
| log_density:nutrientmedium | −1.535 (1.305) | 0.580 (0.481) |
| log_density:nutrienthigh | 0.228 (1.194) | 0.927 (0.572) |
| temperature:log_density:nutrientmedium | 0.108 (0.090) | −0.028 (0.030) |
| temperature:log_density:nutrienthigh | −0.011 (0.082) | −0.056 (0.037) |
| Constant | −8.791 (6.404) | −6.246*** (2.238) |
| Observations | 202 | 205 |
| $R^2$ | 0.529 | 0.546 |
| Adjusted $R^2$ | 0.499 | 0.518 |
| Residual Std. Error | 0.277 (df = 189) | 0.280 (df = 192) |
| F Statistic | 17.669*** (df = 12; 189) | 19.273*** (df = 12; 192) |
| *Note:* | | *p<0.1; **p<0.05; ***p<0.01 |

Table S3

| | *Dependent variable:* | |
|---|---|---|
| | log(MCV) | |
| | Linear | Fast |
| | (1) | (2) |
| temperature | −0.077 (0.065) | 0.142* (0.080) |
| log_density | −0.199 (0.266) | 0.830* (0.448) |
| nutrientmedium | −5.649*** (1.857) | 3.401* (1.926) |
| nutrienthigh | −4.600 (2.853) | 4.068** (1.854) |
| days | | −0.105** (0.052) |
| temperature:log_density | 0.002 (0.014) | −0.041** (0.019) |
| temperature:nutrientmedium | 0.278*** (0.106) | −0.124 (0.080) |
| temperature:nutrienthigh | 0.288* (0.172) | −0.191** (0.078) |
| log_density:nutrientmedium | 0.763** (0.348) | −1.504*** (0.539) |
| log_density:nutrienthigh | 0.545 (0.403) | −1.297** (0.522) |
| temperature:log_density:nutrientmedium | −0.035* (0.019) | 0.063*** (0.023) |
| temperature:log_density:nutrienthigh | −0.030 (0.023) | 0.062*** (0.022) |
| Constant | −4.541*** (1.235) | −8.993*** (1.665) |
| Observations | 205 | 183 |
| $R^2$ | 0.393 | 0.399 |
| Adjusted $R^2$ | 0.358 | 0.357 |
| Residual Std. Error | 0.383 (df = 193) | 0.394 (df = 170) |
| F Statistic | 11.336*** (df = 11; 193) | 9.414*** (df = 12; 170) |

*Note:* *p<0.1; **p<0.05; ***p<0.01

Table S4: Table with SEM coefficients and their standard errors for the different temperature treatments

| | Constant | | Slow | | Linear | | Fast | |
|---|---|---|---|---|---|---|---|---|
| path | estimate | SE | estimate | SE | estimate | SE | estimate | SE |
| Cell size ~bacteria abundance | -0.079 | 0.044 | 0.071 | 0.054 | 0.003 | 0.068 | 0.061 | 0.109 |
| Consumer abundance ~bacteria abundance | -0.492 | 0.178 | -0.091 | 0.2 | -0.063 | 0.221 | -1.044 | 0.261 |
| Oxygen content ~bacteria abundance | 0.629 | 0.194 | -0.527 | 0.2 | 0.099 | 0.18 | 0.534 | 0.192 |
| Growth rate ~bacteria abundance | -0.087 | 0.043 | 0.07 | 0.046 | 0.167 | 0.036 | -0.066 | 0.048 |
| Consumer abundance ~Growth rate | -2.811 | 0.27 | -2.402 | 0.283 | -1.869 | 0.396 | -0.482 | 0.397 |
| Cell size ~Growth rate | 0.211 | 0.078 | 0.246 | 0.086 | -0.07 | 0.128 | -0.184 | 0.16 |
| Oxygen content ~temperature | 0.129 | 0.173 | 0.082 | 0.018 | 0.044 | 0.018 | -0.04 | 0.021 |
| Growth rate ~temperature | -0.001 | 0.037 | -0.041 | 0.004 | -0.066 | 0.003 | -0.126 | 0.005 |
| Consumer abundance ~temperature | -0.048 | 0.153 | 0.049 | 0.021 | 0.031 | 0.033 | 0.036 | 0.058 |
| Cell size ~temperature | -0.033 | 0.036 | -0.008 | 0.006 | -0.041 | 0.01 | -0.111 | 0.023 |
| Bacteria abundance ~temperature | -0.085 | 0.056 | 0.018 | 0.006 | 0.029 | 0.006 | 0.021 | 0.008 |
| Cell size ~Consumer abundance | -0.122 | 0.016 | -0.085 | 0.017 | -0.049 | 0.021 | 0.143 | 0.029 |
| bacteria abundance ~nutrients | -0.152 | 0.035 | -0.093 | 0.029 | -0.086 | 0.033 | -0.151 | 0.037 |
| Cell size ~nutrients | 0.041 | 0.033 | 0.088 | 0.036 | 0.119 | 0.05 | -0.043 | 0.06 |
| Oxygen content ~nutrients | -1.682 | 0.049 | -1.682 | 0.049 | -1.682 | 0.049 | -1.682 | 0.049 |
| Consumer abundance ~nutrients | 1.013 | 0.101 | 1.142 | 0.097 | 1.297 | 0.126 | 0.549 | 0.14 |
| Growth rate ~nutrients | 0.101 | 0.024 | 0.13 | 0.021 | 0.169 | 0.018 | 0.047 | 0.025 |
| Cell size ~Oxygen content | -0.068 | 0.01 | -0.068 | 0.01 | -0.068 | 0.01 | -0.068 | 0.01 |

Table S5: Spearman's rank correlation coefficients ($rho$) between aspect ratio (AR) and cell size (MCV) and corresponding p-values by treatment.

| Temperature | Nutrient | Rho | p-value |
|---|---|---|---|
| constant | low | -0.82 | 5.894670e-20 |
| constant | medium | -0.60 | 5.025890e-09 |
| constant | high | -0.70 | 2.873411e-12 |
| fast | low | 0.14 | 3.056213e-01 |
| fast | medium | -0.72 | 1.886859e-12 |
| fast | high | -0.48 | 4.441602e-05 |
| linear | low | 0.09 | 4.628842e-01 |
| linear | medium | -0.77 | 1.575486e-15 |
| linear | high | -0.58 | 3.752201e-08 |
| slow | low | -0.38 | 5.730952e-04 |
| slow | medium | -0.61 | 3.105849e-09 |
| slow | high | -0.85 | 5.979670e-23 |

#### 2 Supplementary figures

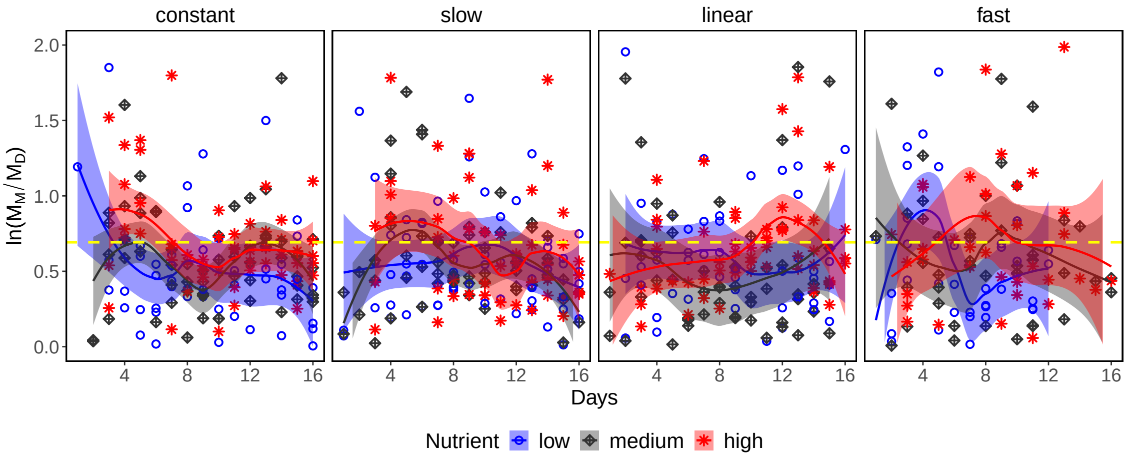


Figure S1: **The ratio of mother and daughter cell sizes of Colpidium striatum**. Cell sizes were corrected for the effect of population abundance. The horizontal yellow dashed line depicts the expected ratio of mother and daughter cell sizes ($ln(2)$).

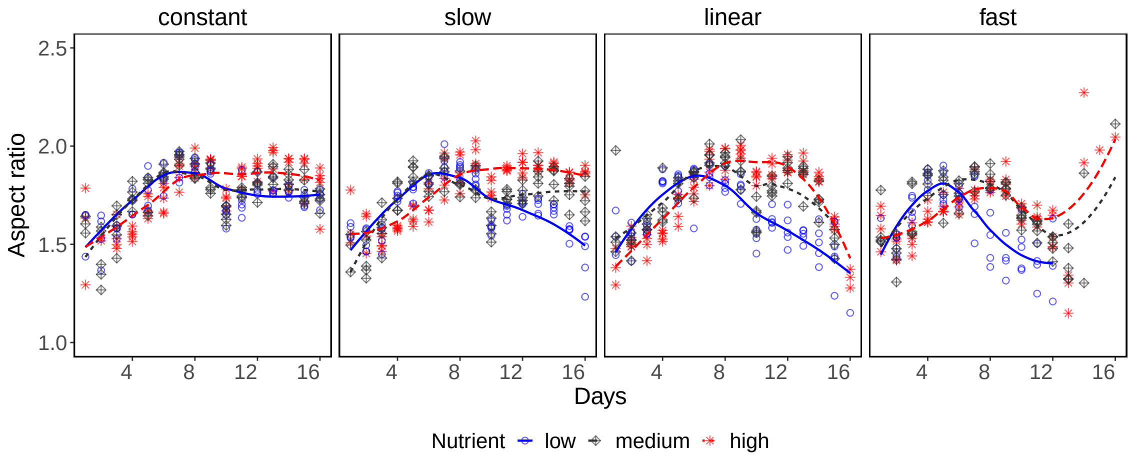


Figure S2: **Aspect ratio of Colpidium striatum**. Points represent the median aspect ratio of cells of each community per day.

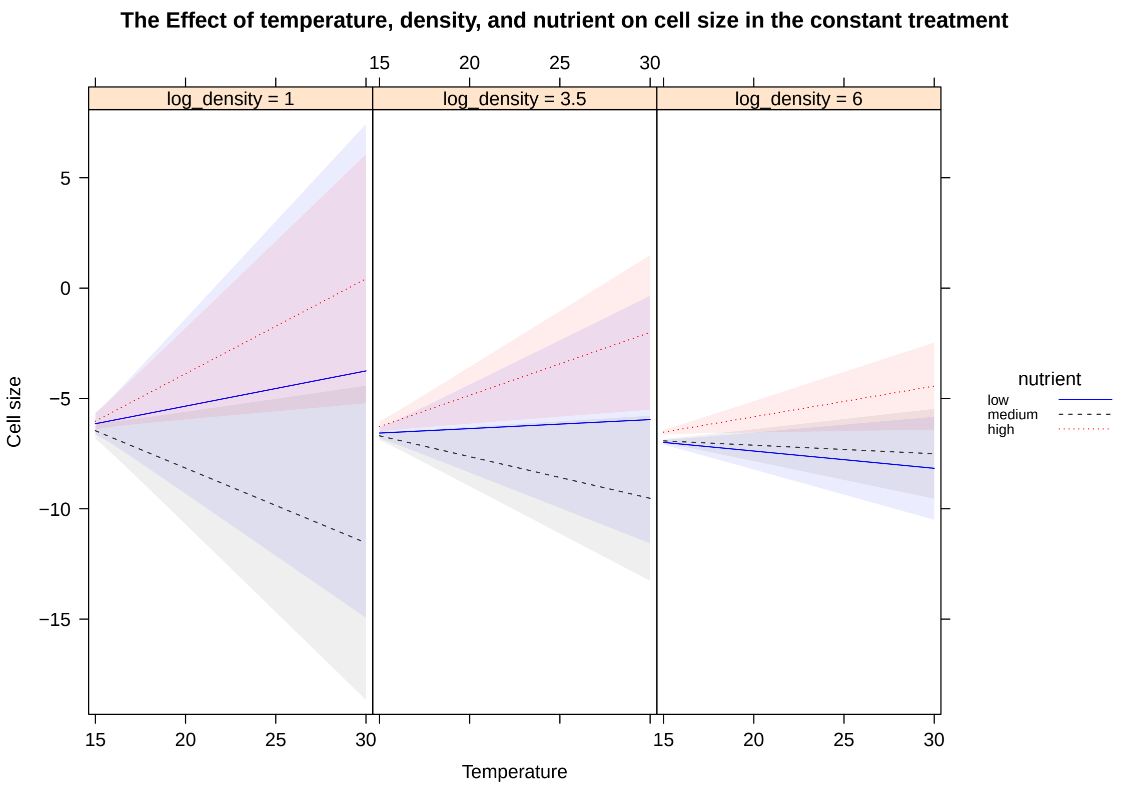


Figure S3: **Three-way interactions in the constant temperature treatment.**

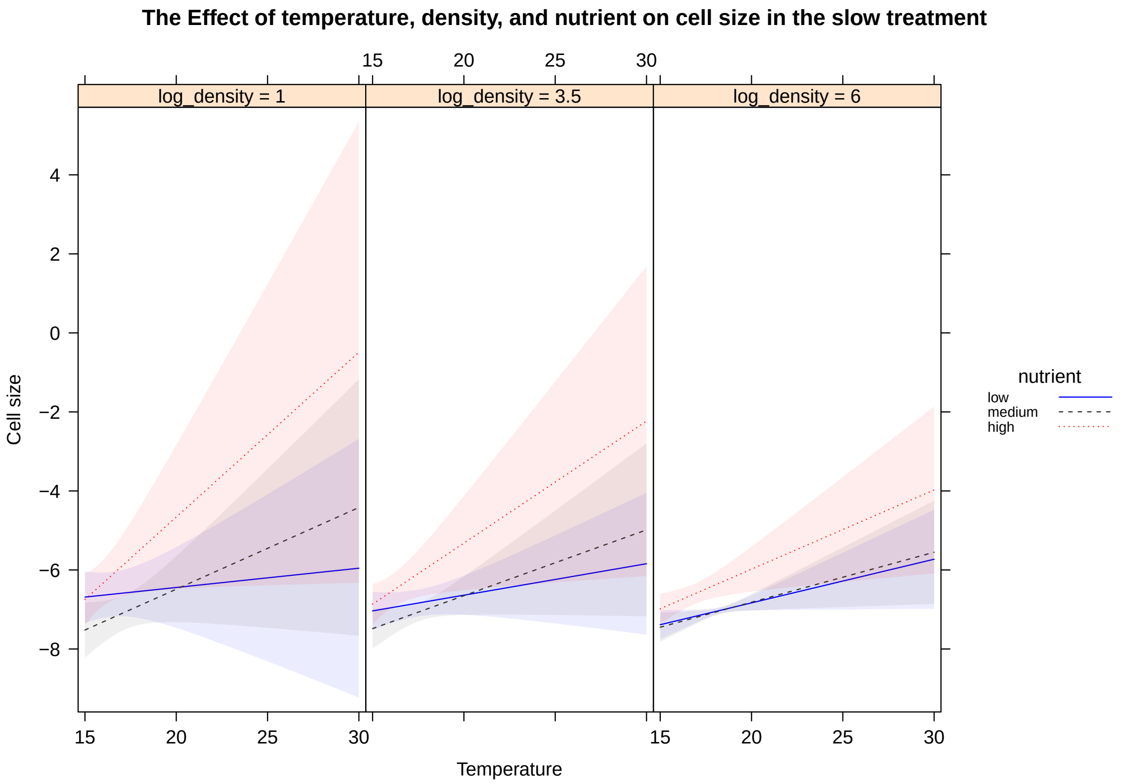


Figure S4: **Three-way interactions in the slow temperature treatment.**

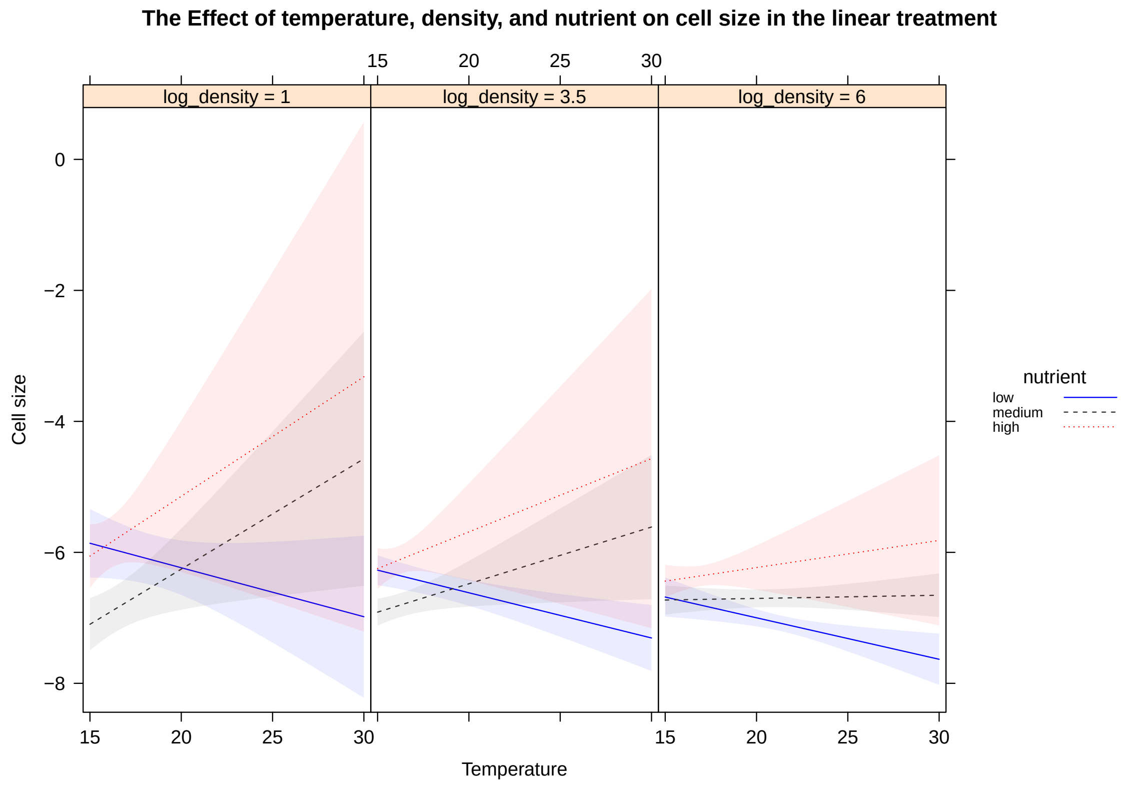


Figure S5: **Three-way interactions in the linear temperature treatment.**

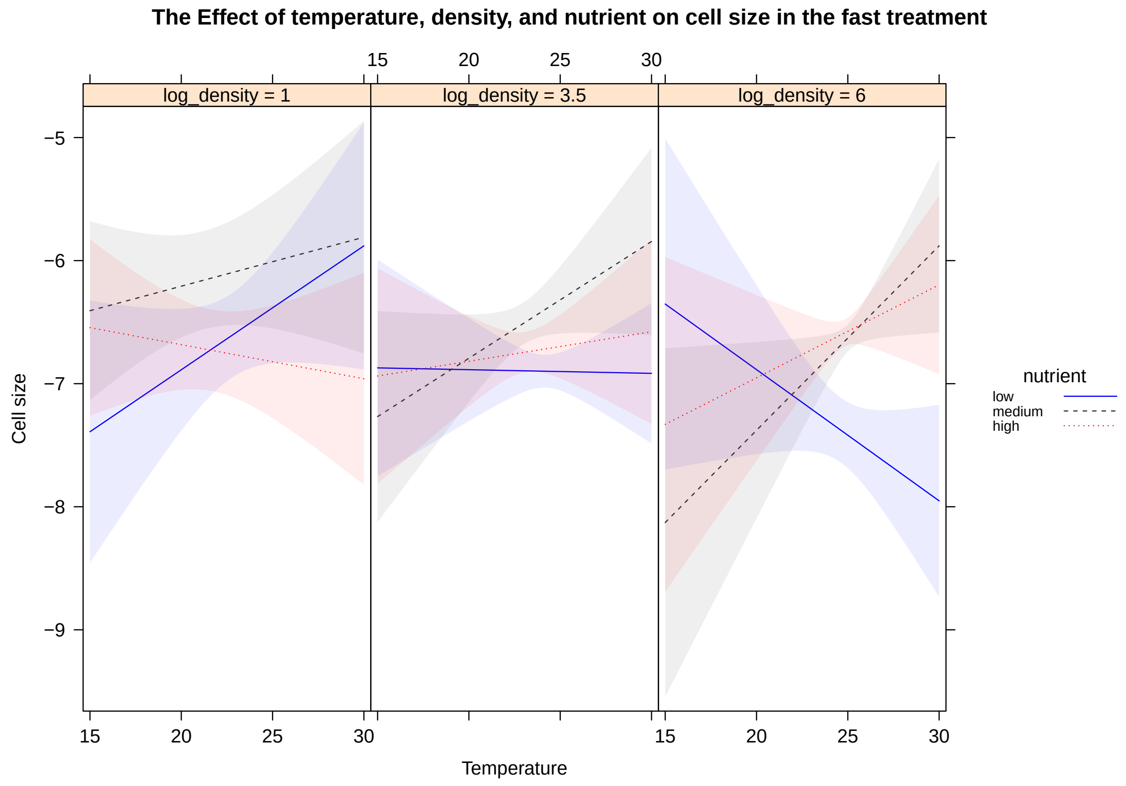


Figure S6: **Three-way interactions in the fast temperature treatment.**
